## Supplementary figures and images for "An unscheduled switch to endocycles induces a reversible senescent arrest that impairs growth of the *Drosophila* wing disc"

### Fig. S1

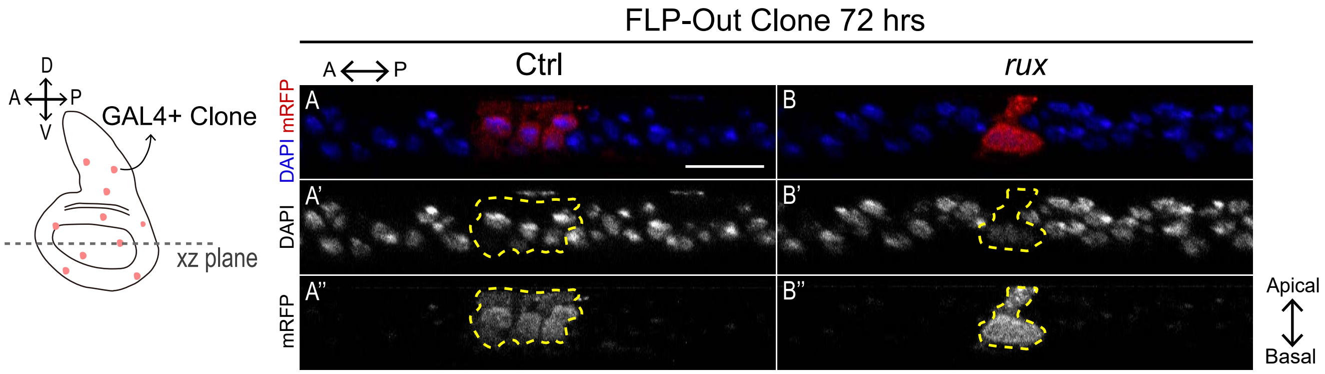

### Fig. S2

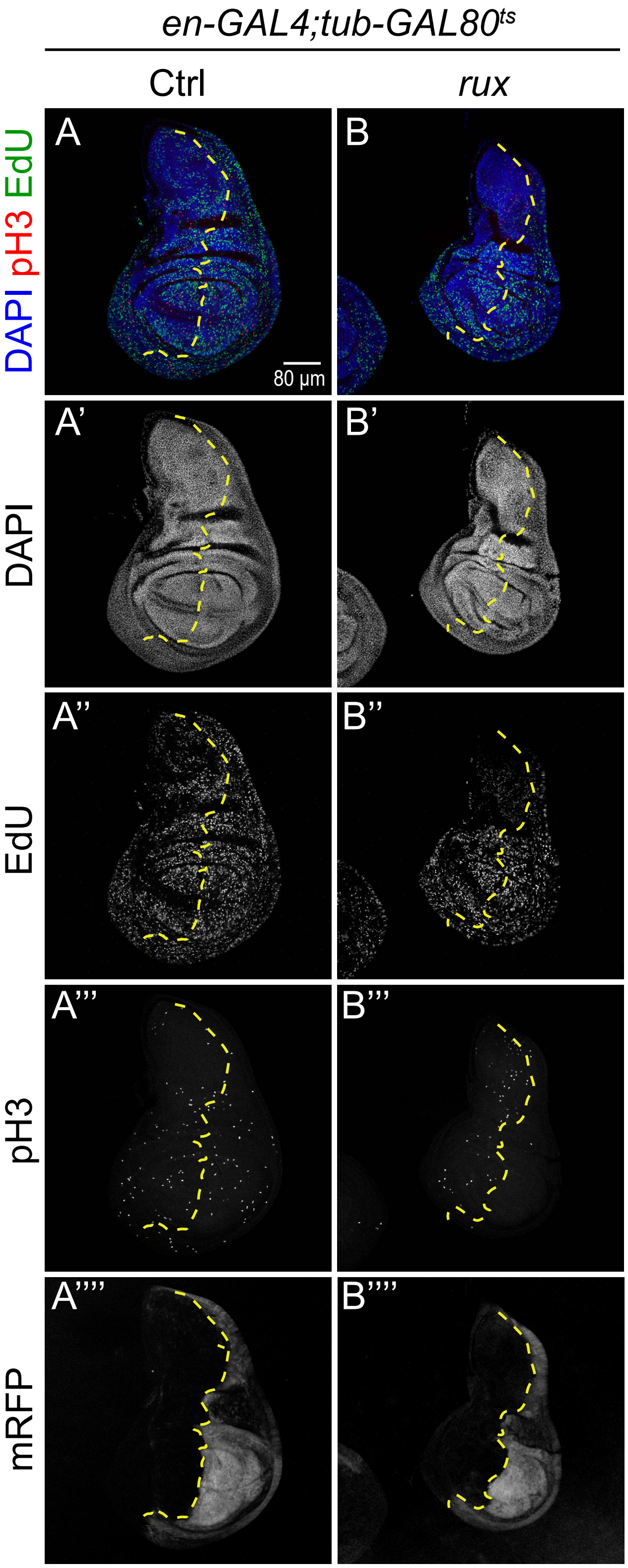

### Fig. S3

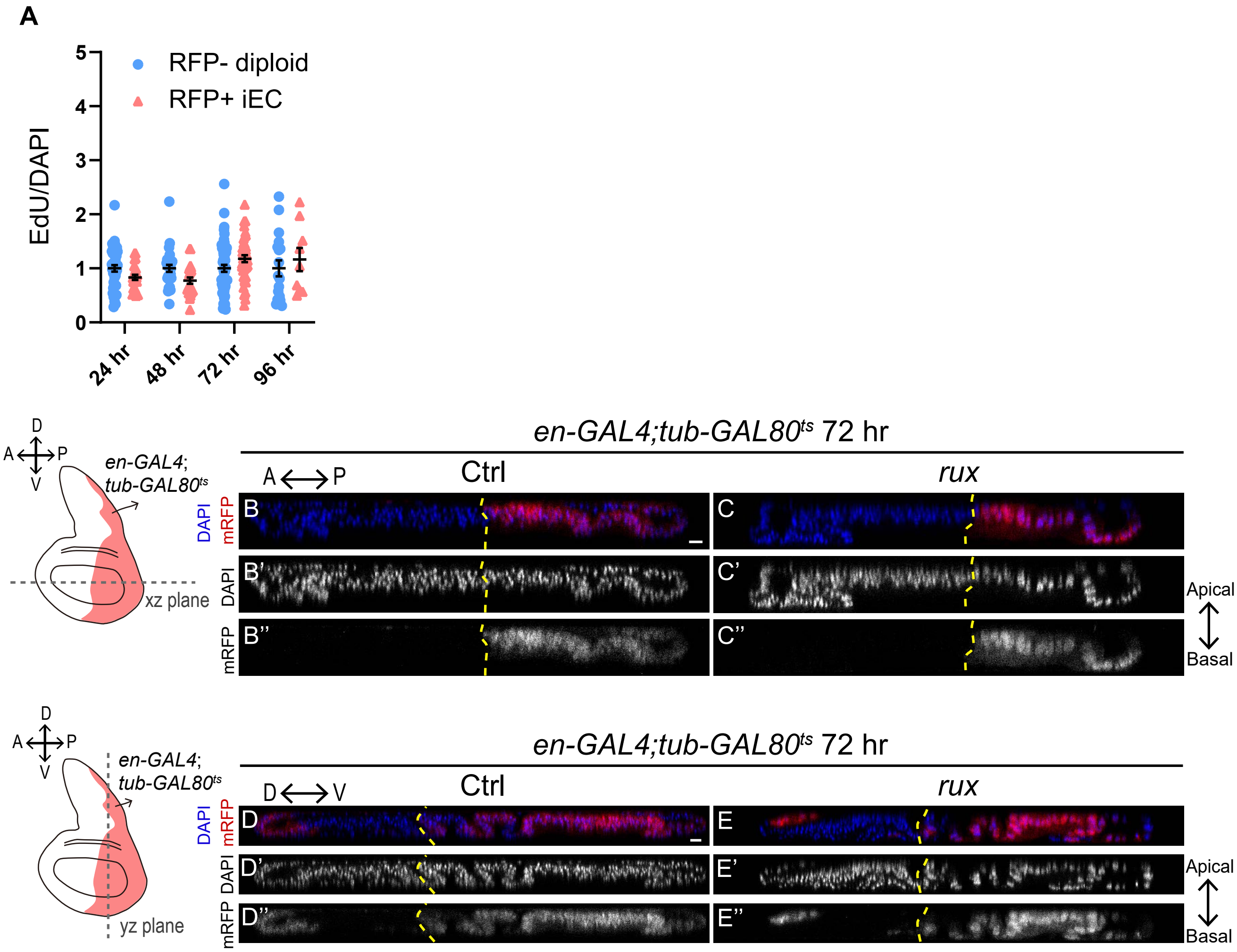

### Fig. S4

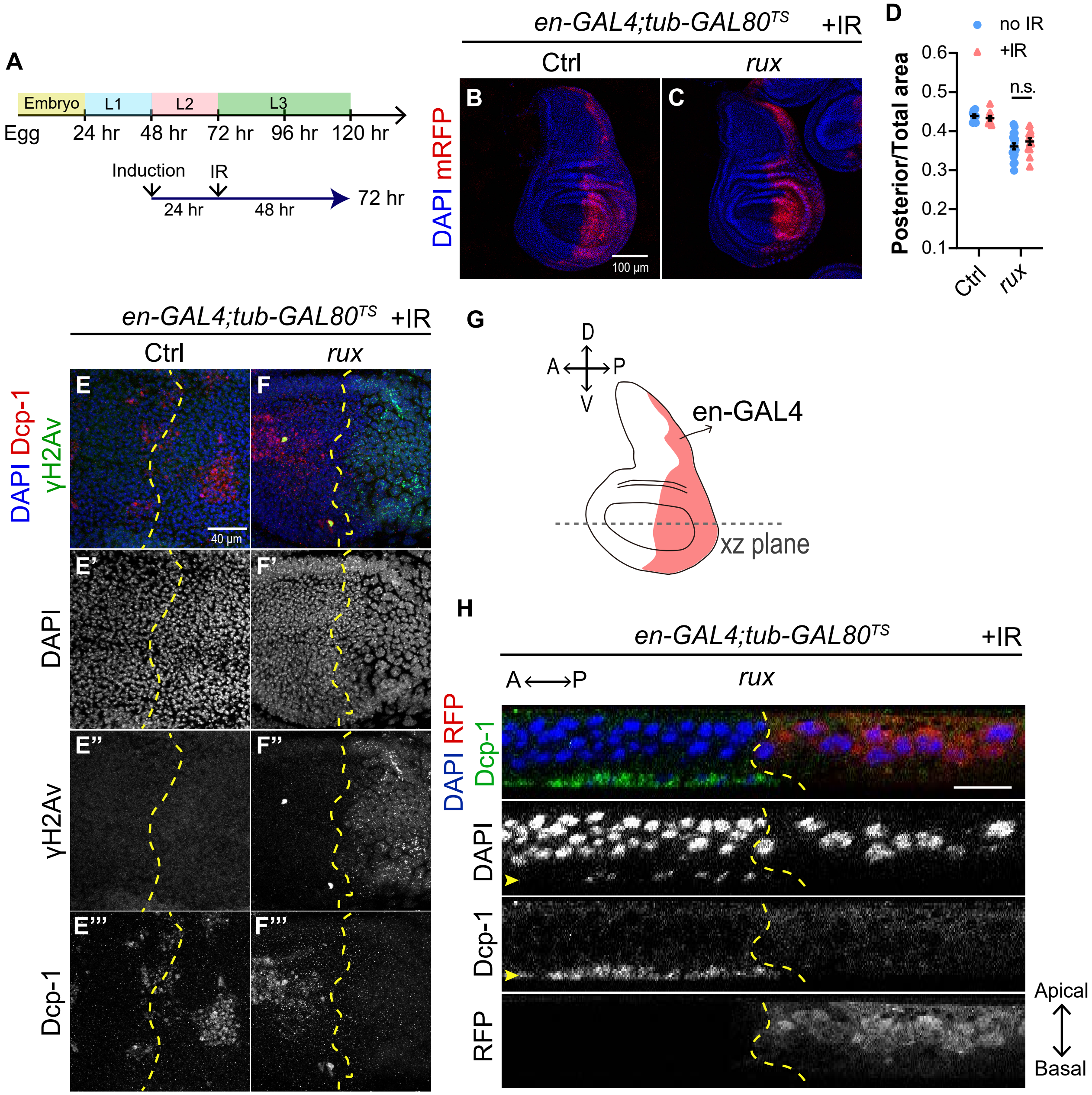

### Fig. S5

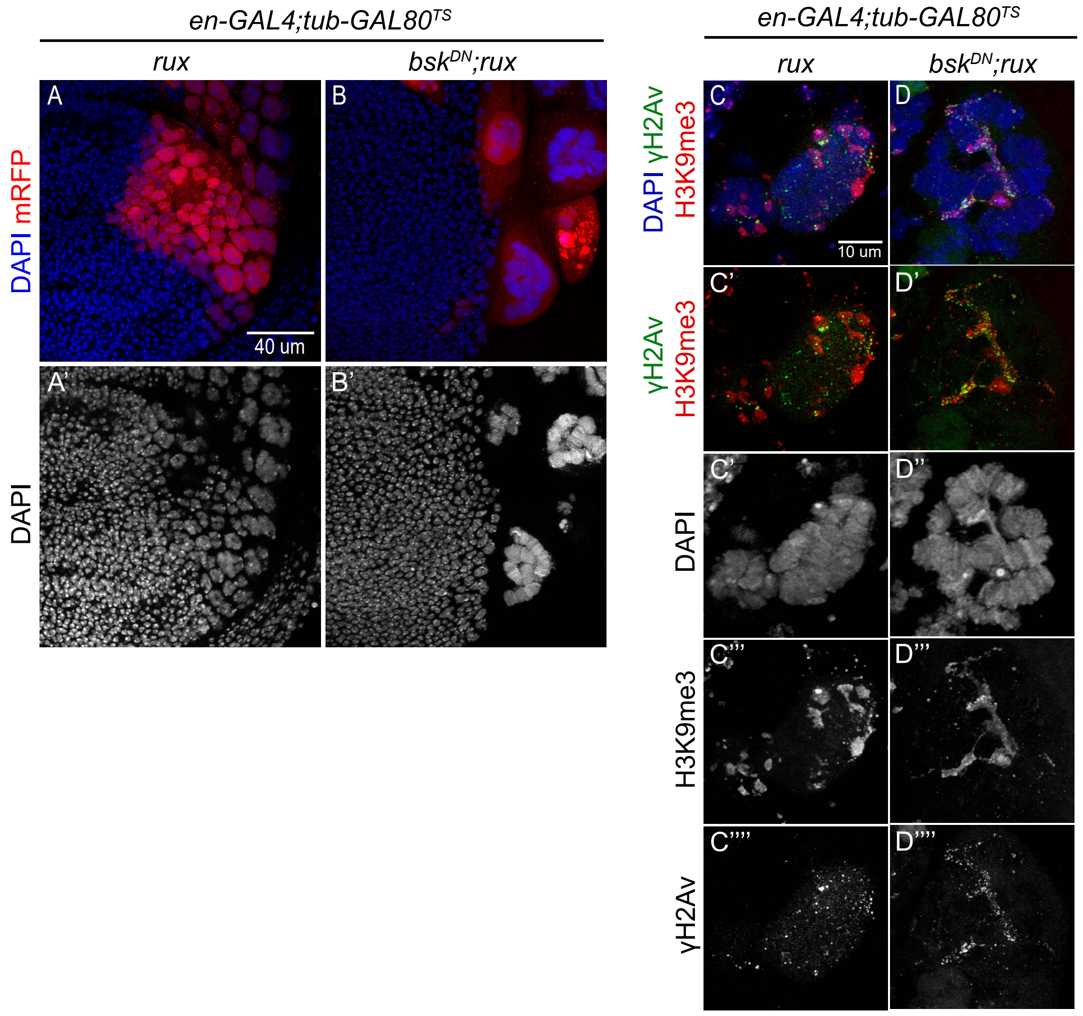

### Fig. S6

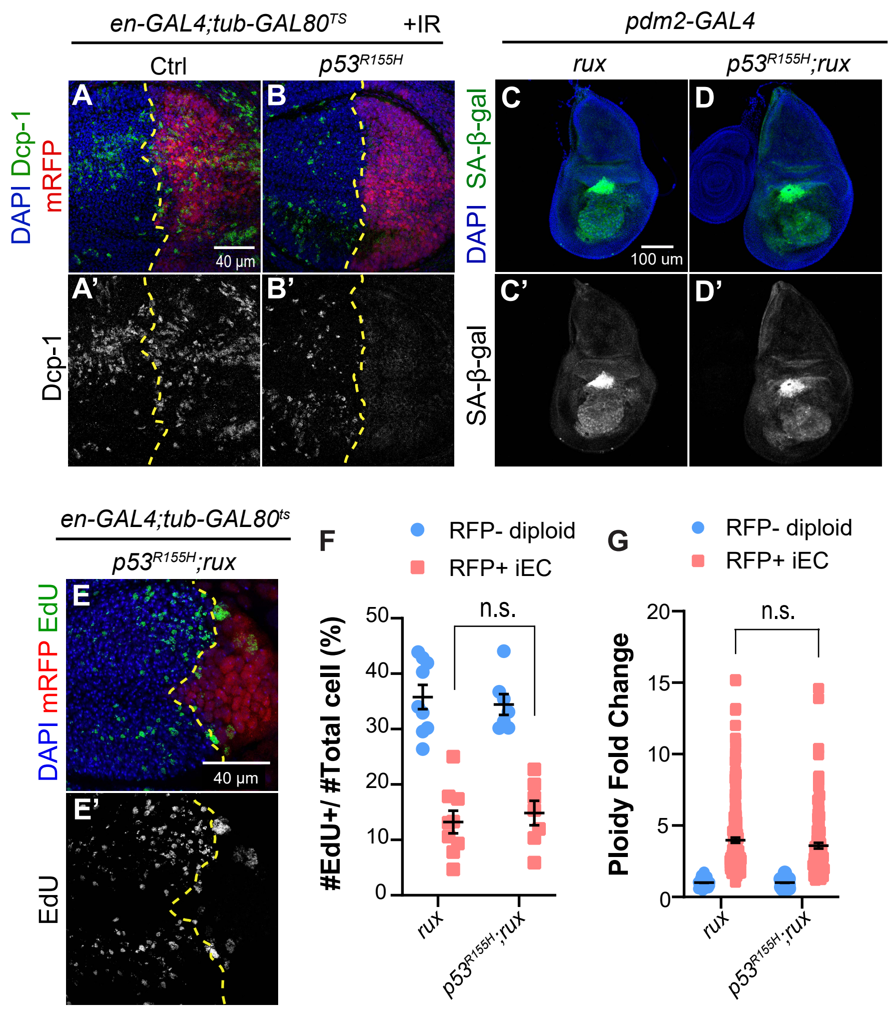
